# Longitudinal Variability of *Bifidobacterium* Species in the Infant Gut is Independent of Maternal Milk HMO Composition

**DOI:** 10.1101/2023.07.17.549354

**Authors:** Dena Ennis, Shimrit Shmorak, Evelyn Jantscher-Krenn, Moran Yassour

## Abstract

The development of the infant gut microbiome is primarily influenced by the infant feeding type, with breast milk serving as the optimal source of nutrition. Breast milk contains human milk oligosaccharides (HMOs) that act as nourishment for the developing gut microbiome, potentially conferring advantages to specific bacterial species. Previous studies have demonstrated the ability of certain *Bifidobacterium* species to utilize individual HMOs, however a longitudinal study examining the evolving microbial community at a high resolution in the context of mothers’ milk HMO composition is lacking. Here, we explored the relationship between the HMO composition in mothers’ milk and the abundance of *Bifidobacterium* species in the infant gut throughout the course of early life. To enable subspecies taxonomic classification, we developed a high-throughput method for quantifying the abundance of *Bifidobacterium longum* subsp. *infantis* (*BL. infantis*; the best known HMO-utilizer) from metagenomic sequencing. We applied this method to a longitudinal cohort consisting of 21 mother-infant dyads, from whom we collected matched breast milk and infant stool samples at multiple time points during the first year of life. We observed substantial changes in the infant gut microbiome over the course of several months, while the HMO composition in mothers’ milk remained relatively stable. *Bifidobacterium* species were a prominent factor contributing to the variation observed among samples; however, no significant associations were found between specific HMOs in mothers’ milk and the abundance of *Bifidobacterium* species. Finally, the longitudinal nature of our cohort enabled us to characterize the dynamic colonization of *BL. infantis* in the infant gut, which surprisingly began late in the breastfeeding period. Applying our *BL. infantis* quantification method to additional datasets from various geographical locations, we found similar, late-colonization by *BL. infantis*, highlighting the importance of quantifying *BL. infantis* in the infant gut.

## Introduction

Breast milk is considered the ideal nutrition for infants during their first six months of life^1^. Millions of years of evolution have shaped breast milk composition such that its third most abundant component, human milk oligosaccharides (HMOs), cannot be digested by the infant, but serves as substrate for the infant’s gut bacteria^2, 3^. There are many different types of HMOs, which can be largely classified into three groups: fucosylated, sialylated, or neutral. Each HMO is composed of 3 to 32 monomers, and a single milk sample typically contains 50 to 200 distinct types of HMOs^4^. Among other factors, maternal genetics plays a role in the production of specific HMOs in breast milk^5^. For example, mothers with an inactive fucosyltransferase 2 (*FUT2*) gene, termed non-secretors, fail to form alpha-1,2 bonds between fucose and lactose or other HMO backbone structures, resulting in the lack of 2’FL and other alpha-1,2-fucosylated glycans^6^. Additionally, environmental factors coupled with infant age can affect HMO composition^7^.

On average, infants have a higher relative abundance of *Bifidobacterium* species while they are breastfed^8–10^. *Bifidobacterium* species were previously shown capable of utilizing multiple HMOs^11–13^, however this ability varies between species and even within a single species^14–17^. Among all *Bifidobacterium* species, the best-known HMO utilizer is *Bifidobacterium longum* subsp*. infantis* (*BL. infantis*) which grows efficiently on most types of HMOs^18^, and possesses a large variability of HMO utilizing genes^19^. In contrast, other *Bifidobacterium* species have a lower capability of HMO utilization, for example *B. breve* strains cannot utilize 3’SL and 6’SL at all, and most of them cannot utilize fucosylated HMOs^15^. Since HMOs serve as food for the gut microbiome, one may hypothesize that different HMO compositions in mothers’ milk affects the developing gut microbial community.

To date, most research addressing the HMO-bacteria relationship in the infant gut focused on a single time point^20^. A longitudinal cohort study is needed in order to examine how changes in HMO composition impact the infant gut microbiome over time.

To quantify the abundance of various *Bifidobacterium* species in microbiome communities two approaches are commonly used: 16S-rRNA sequencing and shotgun metagenomics. While metagenomics allows classification at the species level, 16S-rRNA sequencing provides only genus-level classification of microbiome communities. The basic annotated unit in 16S-rRNA sequencing is referred to as operational taxonomic unit (OTU), which can be assigned to genus-level classification and may represent an unknown sub-clade of the assigned genus. Multiple species can be annotated as the same OTU, hence using 16S-rRNA sequencing so far provided mostly weak or no associations with abundances of specific HMOs ^21–24^. A single OTU can include multiple species (or subspecies) with various HMO utilization capabilities, thus, a higher-resolution taxonomic definition is needed.

The largest variability in HMO-utilization capability can be found within the *Bifidobacterium longum* species. Overall, this species can be divided into two subspecies found in humans; *B. longum* subsp. *longum* (*BL. longum*) and *B. longum* subsp. *infantis* (*BL. infantis*). *BL. longum* is found both in infants and adults, while *BL. infantis* is unique to the infant gut. Studies have shown that *BL. infantis* can utilize almost all HMOs^25^, while *BL. longum* has a limited repertoire. To study the HMO-microbe relationship, taking into account these differences in HMO-utilization within *B. longum* subspecies, a high-throughput, higher-resolution method is needed. Past studies have used different methods to differentiate between *BL. infantis* and *BL. longum*, such as qPCR^26^, PCR^27^ or the Bifidobacterium Longum-Infantis Ratio (BLIR) method^28, 29^, yet these methods require the original DNA and are not high-throughput. Others have searched for *BL. infantis* specific genes such as the H1 cluster^20^ or other *BL. infantis* clusters^30, 31^, however these methods do not give an exact ratio between the subspecies, rather they indicate their presence or absence. Ideally, the newly developed method could be applied also to the massive amounts of existing data in the literature.

Here, we established a new matched cohort of breast milk and infant stool samples collected longitudinally throughout the first year of life. We developed a method to allow *BL. infantis* quantification from existing metagenomic data, and applied it to samples from our cohort to study the relationship between the abundance of *Bifidobacterium* species in the infant gut and HMO composition in mothers’ milk over time. Finally, we applied our *B. longum* subspecies quantification method to existing infant gut datasets to examine differences in *BL. infantis* prevalence across geographic locations.

## Results

### Cohort design

We have established a new and unique longitudinal cohort to test the relationship between HMOs in mothers’ milk and the developing infant gut microbiome. Our cohort consists of 21 mother-infant dyads with 83 matched infant stool and breast milk samples collected from the same day, together with the infant nutritional information and antibiotic treatments **(Supplementary Figure 1, Supplementary Table 1)**. We collected these samples between the age of two weeks and 41 weeks, and each dyad contributed between one to eight samples.

### Specific marker genes allow better quantification of *B. longum* subspecies

*Bifidobacterium longum subsp. infantis* (*BL. infantis*) is the best known utilizer of HMOs^19, 32^, however current methods for metagenomic classification are unable to separate the *Bifidobacterium longum* (*B. longum*) species into its subspecies; *Bifidobacterium longum subsp. longum* (*BL. longum*) and *BL. infantis*^20, 29^. MetaPhlAn is one of the most common tools for profiling the composition of microbial population from metagenomic data, by using specific marker genes for each taxonomic group^33^. However MetaPhlAn has no specific markers for *BL. infantis* and therefore classifies *B. longum* at the species taxonomic level. Due to the differences between *B. longum* subspecies in the context of HMO utilization, there is a rising need for a high-throughput method that will allow specific identification and quantification of *BL. infantis* from metagenomics data.

Here we define *B. longum* subspecies specific markers and use them in a tailored MetaPhlAn^33^ database which allows abundance quantification of *B. longum* subspecies **(Supplementary Figure 2A)**. To construct our new dataset, we searched for marker genes that are unique to each subspecies. A marker gene was selected if it was present in 90% of reference genomes of one subspecies and not in a single genome of the other subspecies (**Figure 1A, Methods)**. The variation of gene presence differed within the two subspecies; while *BL. infantis* displayed a tightly defined taxonomic group, *BL. longum* showed higher diversity across genomes (**Figure 1A**). The difference can be attributed to the specific definition of *BL. infantis* strains, which are isolated from infants and possess gene clusters facilitating HMO utilization^14, 19^, whereas *BL. longum* encompasses a broader range of strains referred to as “all other” strains.

**Figure 1:**
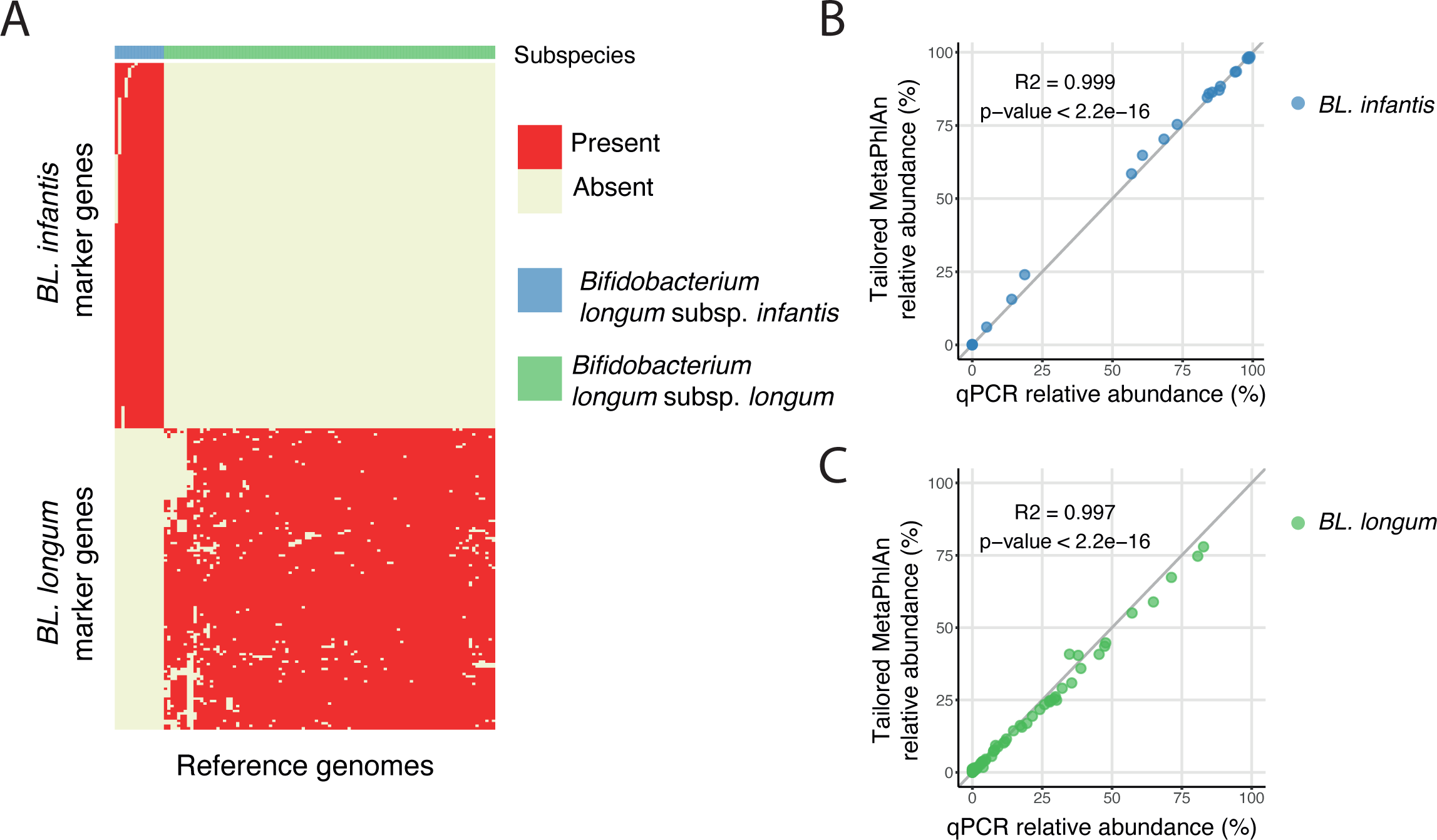
Quantification of *Bifidobacterium longum* subspecies using marker-genes enables subspecies-level detection. **(A)** Identification of unique genes in *BL. infantis* (blue) and *BL. longum* (green) reference genomes that can serve as marker-genes in the quantification of *B. longum* subspecies. **(B-C)** Validation of the computational approach, comparing the tailored MetaPhlAn marker-gene quantification results to the experimental qPCR results of **(B)** *BL. infantis* and **(C)** *BL. longum*.

In order to validate our results, we applied MetaPhlAn with our new set of marker genes coupled with subspecies-specific qPCR to metagenomic sequencing data from 68 infant stool samples. When comparing the relative abundance of *BL. infantis* and *BL. longum* in each method, we observed a strong correlation between our computational approach and qPCR (R^2^=0.999 for *BL. infantis* and 0.997 for *BL. longum*; **Figure 1B,C**). This finding confirms our method’s specificity and sensitivity for both *BL. infantis* (**Figure 1B**) and *BL. longum* (**Figure 1C**). In some samples, MetaPhlAn failed to assign a classification to a small percentage of *BL. longum* and therefore it was designated as unclassified *B. longum* **(Supplementary Figure 2B,C)**. This observation could potentially be attributed to the extensive diversity within the *BL. longum* clade.

### The infant gut microbiome shows excessive changes while HMO composition in mothers’ milk is fairly stable

To examine the dynamics of *Bifidobacterium* species in the infant gut, we conducted metagenomic sequencing and analyzed the data using our novel MetaPhlAn database. We observed a significant prevalence of *Bifidobacterium* (at least one sample with >25%) in all the infants (**Figure 2A**), in line with our expectation as most infants in our study were breastfed^8, 14, 34^. *Bifidobacterium* remained highly abundant even after solid foods were introduced to infants (**Figure 2A**, arrows). The abundant species included *Bifidobacterium breve*, *Bifidobacterium bifidum*, *Bifidobacterium pseudocatenulatum*, *B. longum* subspecies (**Figure 2A**), along with *Bacteroides* species such as *Bacteroides dorei* and *Bacteroides vulgatus* **(Supplementary Figure 3)**. Interestingly, the presence of *BL. longum* and *BL. infantis* was mutually exclusive, reflecting potential intra-species competition, as previously suggested^35^.

**Figure 2:**
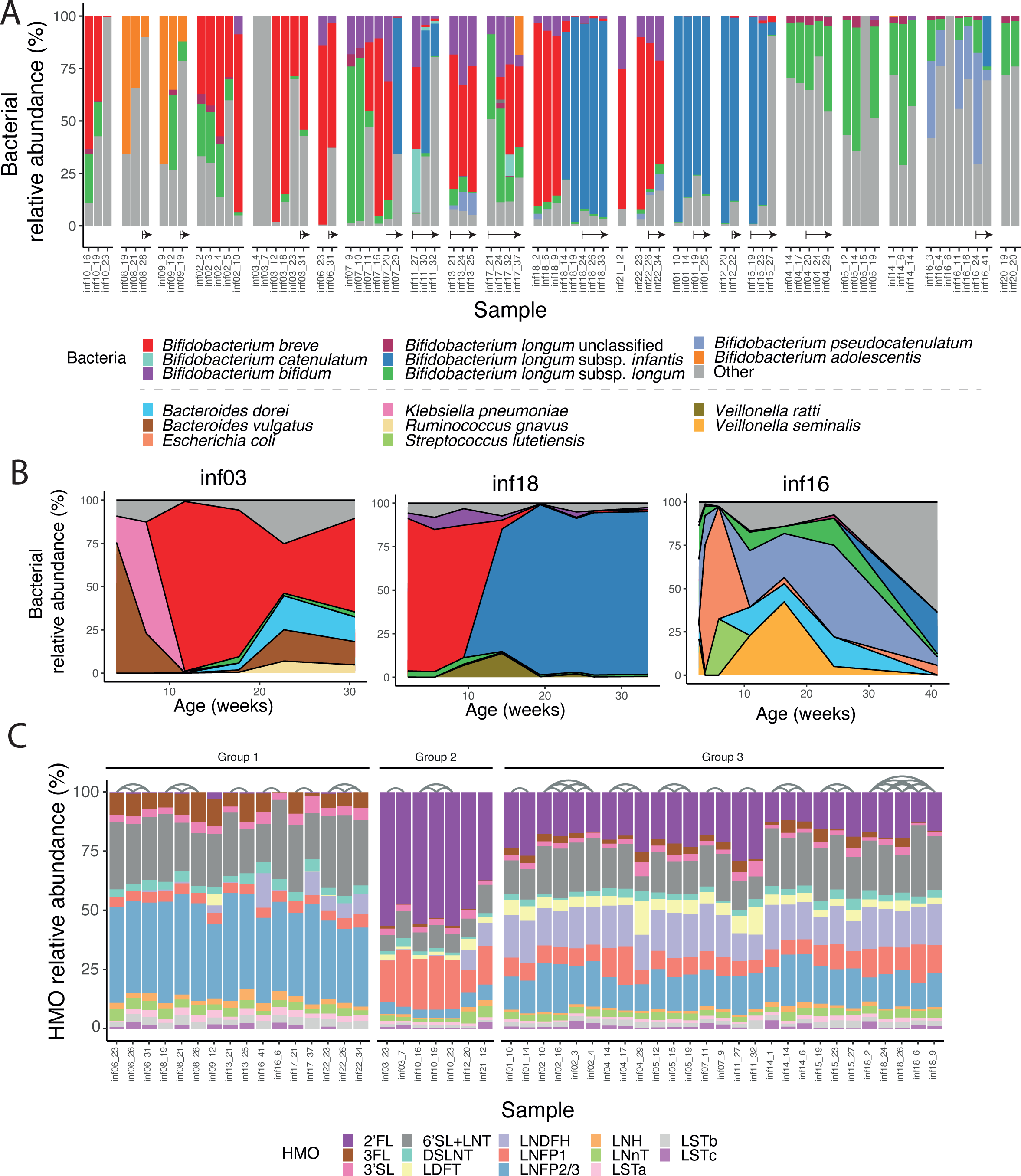
Composition and dynamics of *Bifidobacterium* species in the infant gut and HMOs in mothers’ milk. **(A)** Relative abundance of *Bifidobacterium* species in the infant gut (colorful) with all other species classified as “other” (gray). Samples from the same infant are grouped together, sorted by age. Arrows indicate samples taken after the introduction of solid food. **(B)** Temporal changes in the relative abundance of the microbial community in three infants (inf03, inf16, and inf18) in the first 30-40 weeks of life. The most prevalent bacteria are colored, and the remaining are indicated as “Other” (gray). Colors as in (A), with additional colors for the non-*Bifidobacterium* species. **(C)** Relative abundance of 16 HMOs measured in mothers’ milk. Samples are categorized into three groups based on their HMO profiles, and arches connect samples obtained from the same mother (**Methods**).

Overall, we found that the infant gut microbiome underwent significant changes over the course of several months. While the bacterial composition tended to be stable over a few weeks, there were certain time points when a switch in composition occurred (**Figure 2A,B**). In infants that we had samples from many time points we found this switch to occur around ∼10 weeks of life. For example, in infant03 over the course of the first 15 weeks the dominant bacteria in the gut constantly changed, including *B. vulgatus*, *Klebsiella pneumoniae*, and finally *B. breve*; the gut of infant18 was initially dominated by *B. breve*, followed by a complete switch to *BL. infantis*; and the gut of infant16 constantly changed its microbial composition (dominated by *Escherichia coli*, *Veillonella seminalis* and *B. pseudocatenulatum*; **Figure 2B**). It was not always clear what triggered these microbial shifts, however since *Bifidobacterium* and other gut microbes utilize HMOs, we hypothesized that changes in the HMO composition in mothers’ milk might be causing bacterial changes in the infant gut.

To examine the impact of HMO composition in mothers’ milk on the infant gut microbiome, we quantified 16 common HMOs in 50 milk samples from 20 mothers using high performance liquid chromatography with fluorescence detection (HPLC-FLD; **Methods**). In contrast to the dynamic infant gut microbiome, the composition of HMOs in mothers’ milk remained relatively stable over the course of months, in terms of both their concentration in milk **(Supplementary Figure 4A)** and the relative abundance of specific HMOs (**Figure 2C**). We divided the milk samples into three main groups, based on their HMO composition: those with low or no 2’FL abundance (Group 1, non-secretors^36^); samples from secretor mothers with very high abundance of 2’FL (>40%) and LNFP1 (>10%) and smaller amounts of other HMOs (Group 2); and samples from secretor mothers with lower abundance of 2’FL(<30%), and no clear dominant HMO (Group 3; **Figure 2C**). Overall, we found no major changes over time in the abundance of specific HMOs, other than LSTc which was reduced to almost 0 over the course of ∼40 weeks **(Supplementary Figure 4B,C**), in line with previous findings^7^.

### The dominant *Bifidobacterium* species shape the infant gut microbiome

General comparison of the infant gut microbiome composition determined that *Bifidobacterium* species play a significant role in the breastfed infant gut (**Figure 2A**). We next searched for differences across samples in an attempt to characterize the various microbial profiles of the infant gut. We examined the diversity of the infant gut samples in our cohort, using a dimension reduction approach (PCoA with Bray-Curtis dissimilarity, **Methods**), and found that samples cluster into distinct clusters (using K-means, k=4). The first three groups had samples with mostly one main *Bifidobacterium* dominant species (with relative abundance of more than 30%): *B. breve*, *BL. longum* and *BL. infantis*, and the fourth group contained samples dominated by either a different *Bifidobacterium* species (*Bifidobacterium adolescentis*, *B. pseudocatenulatum*) or other species (named “Mixed”, **Figure 3A**). While usually consecutive samples from the same infant were assigned to the same cluster, occasional cluster switches were observed, strengthening our finding that the microbiome changes in this time frame (**Figure 3B**). Notably, when consecutive samples switched groups, there were some enriched transitions, like transitioning into and from the *BL. infantis* group. Overall, these analyses highlight the importance of *Bifidobacterium* species in our samples, as these are major factors that impact the variation in the infant gut microbial composition.

**Figure 3:**
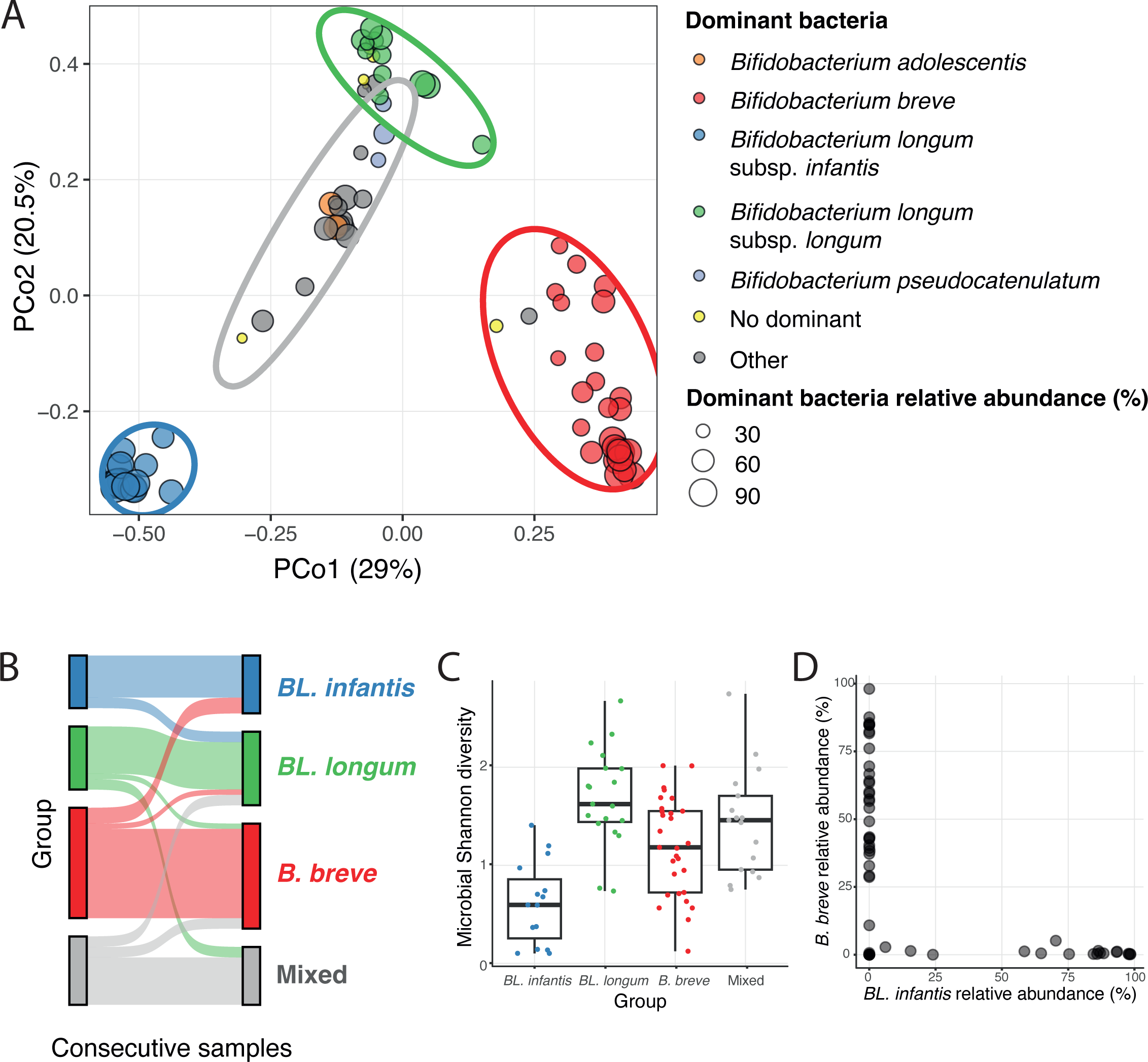
Variation across samples is highly impacted by the dominant *Bifidobacterium* species. **(A)** Principal Coordinate Analysis (PCoA) of infant gut microbiome samples using Bray-Curtis dissimilarity. Points are color-coded based on the dominant *Bifidobacterium* species in each sample, and non-*Bifidobacterium* dominated samples are colored in gray. Samples without a dominant species (>30% relative abundance) are labeled as “No dominant” (yellow). The size of each point represents the relative abundance of the dominant bacteria in that sample. Ellipses encompass four groups identified using k-means clustering. Each group represents a primary dominant species (indicated by the ellipse color): *BL. infantis* (blue), *BL. longum* (green), *B. breve* (red), and “Mixed” (gray). **(B)** Changes in group assignment observed in consecutive samples from the same infant (colors as in A). **(C)** Microbial diversity of samples in each group (measured by the Shannon index; colors as in A). **(D)** Relative abundance of *BL. infantis* (x-axis) versus the relative abundance of *B. breve* (y-axis) in each sample, indicating the mutual exclusiveness of the two species.

To further investigate these four groups of infant gut samples, we examined the microbial diversity of samples found in each group. We found the diversity of samples within the *BL. infantis* group was lower compared to samples from other groups (**Figure 3C**), indicating that the higher abundance of *BL. infantis* leaves a smaller ecological niche for other bacteria in the infant gut. We next focused on the “Mixed” group, and asked whether additional variables may play a role in these microbial profiles. We examined breastfeeding, maternal or infant antibiotic use, delivery mode, breastfeeding type (pumped or direct), and introduction of solid foods, yet we did not find any specific variable that was associated with the microbial profile of the “Mixed” group. As expected, the *BL. infantis* group consisted solely of infants who received none-to-low amounts of infant formula.

To characterize the relationships among dominant *Bifidobacterium* species in the infant gut, we examined their occurrence within groups where they are not dominant. We found that *BL. infantis* and *B. breve* are mutually exclusive, consistent with a previous study in Hazda infants^37^, implying competition for the same niche in the infant gut (**Figure 3D**). However, it remains unclear what specific niche *B. breve* and *BL. infantis* are competing for, given *B. breve*’s limited ability to utilize a variety of HMOs^15^. Finally, we observed *BL. longum* in some *B. breve*-dominant samples (data not shown), suggesting that *B. breve* may rely on derivatives from *BL. Longum* through cross-feeding in these samples^38^.

### Single HMOs are not associated with specific *Bifidobacterium* species

It is well established that different *Bifidobacterium* species have different HMO-utilization capabilities^14, 38, 39^. Therefore, specific HMOs may benefit specific *Bifidobacterium* species in the infant gut based on their HMO utilization profiles. However, we found no significant correlation between the abundance of *Bifidobacterium* in general and the main *Bifidobacterium* species and subspecies with specific HMOs (**Figure 4A**) or HMO groups (fucosylated, sialylated or neutral; **Supplementary Figure 5**). Nevertheless, we found that *BL. infantis* exhibited a high abundance (>25%) exclusively in infants to secretor mothers (**Figure 4B**). In addition, we observed a modest and non-significant negative correlation (r=-0.27) between *BL. infantis* and LSTc (**Figure 4C**). It is worth noting the importance of considering the timing factor in interpreting these findings. The delayed presence of *BL. infantis* in the gut (which will be discussed in more detail later) and the gradual decrease of LSTc over time **(Supplementary Figure 4B,C)** could contribute to the observed correlation. The lack of variation in the HMO composition together with the lack of HMO-microbes associations indicate that the microbial shifts, specifically within *Bifidobacterium* species, can not be explained by a change in mothers’ milk HMO composition.

**Figure 4:**
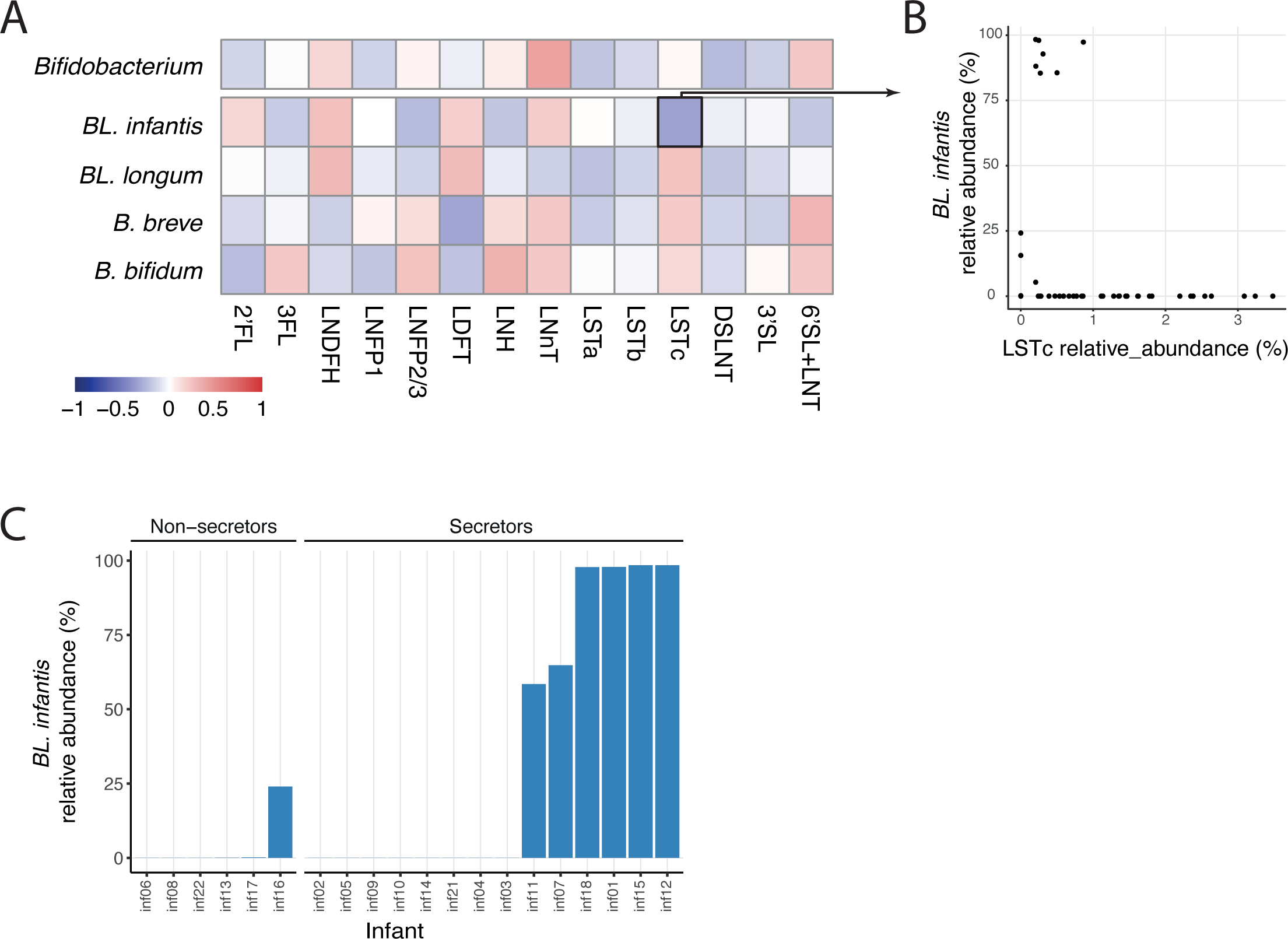
Mothers’ milk HMO composition shows no significant correlation with infant gut *Bifidobacterium*. **(A)** Pearson correlations between all measured HMOs and the *Bifidobacterium* genus, as well as the main individual species and subspecies (*BL. infantis*, *BL. longum*, *B. breve* and *B. bifidum*). None of these correlations were found to be statistically significant. **(B)** Comparison of the relative abundance of LSTc in mothers’ milk (x-axis) with the relative abundance of *BL. infantis* in the infant gut microbiome (y-axis). **(C)** Maximum relative abundance of *BL. infantis* in each infant divided by infants with secretor and non-secretor mothers, highlighting the high abundance of *BL. infantis* only in infants to secretor mothers, but not in all of them.

### Metagenomes with *BL. infantis* contain more HMO utilizing genes

Metagenomes obtained from various time points of multiple infants contain distinct strains and species, resulting in variable gene abundance profiles which can enable various patterns of HMO utilization. To assess the HMO utilization potential of specific *Bifidobacterium* species in our dataset, we investigated the presence of HMO-utilizing genes (HUGs)^40^ organized into five distinct clusters (H1-H5^32^; **Figure 5**). We observed that the dominant species in each sample significantly influenced the metagenome’s theoretical capacity for HMO utilization. As expected, *BL. infantis*-dominated samples exhibited the highest abundance of HUGs, confirming its exceptional capability in utilizing HMOs^19^. Notably, some of these samples displayed high variation in gene abundance from clusters H1 and H5, indicating a potential lower capacity to transport some HMOs^19^, and utilize lacto-N-tetraose (LNT) and lacto-N-neotetraose(LNnT)^16^.

**Figure 5:**
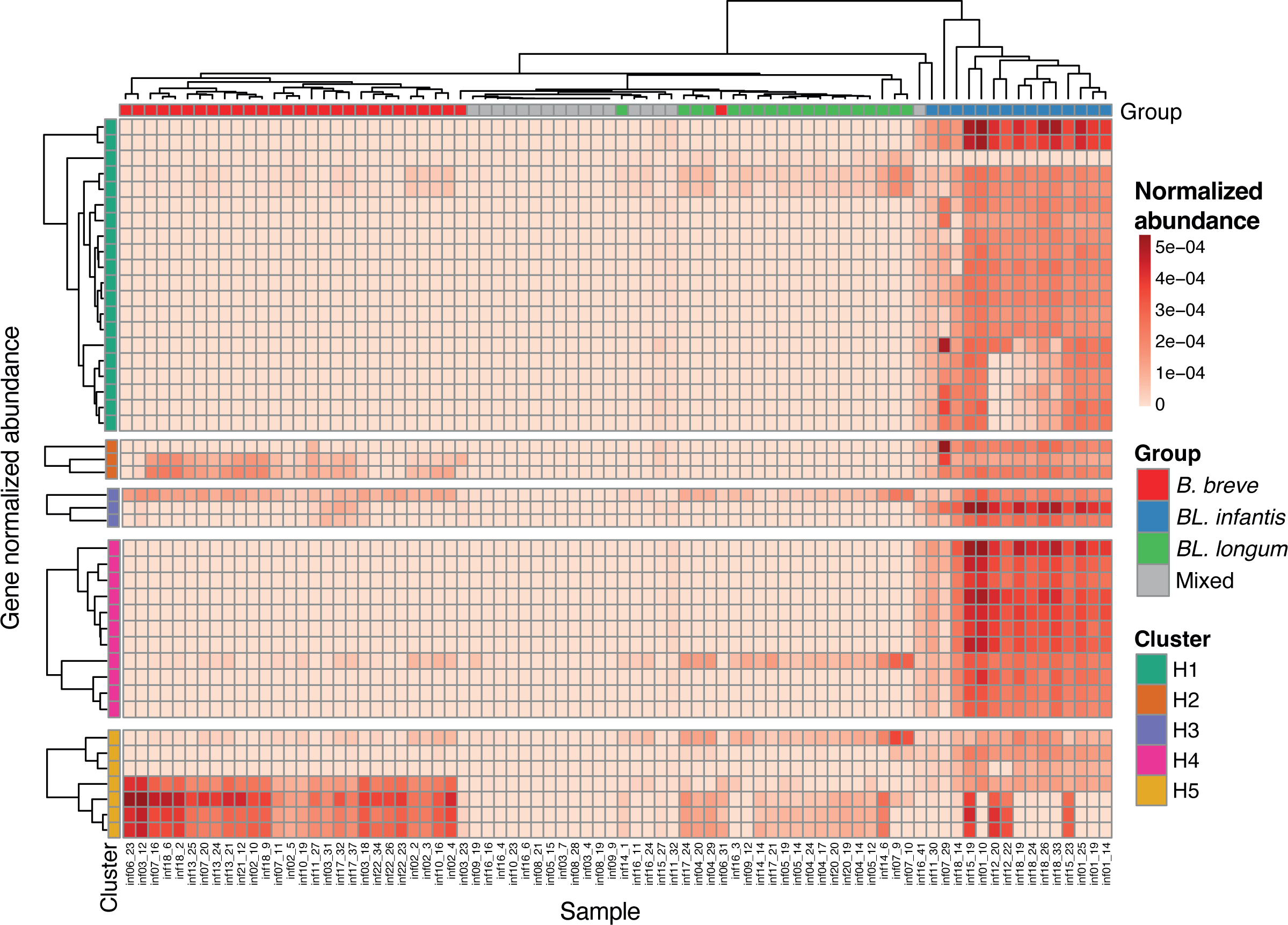
Abundance of HMO utilization genes in metagenomic samples. Normalized genes abundance of HMO utilization genes in all infant samples. The genes are categorized and labeled according to their respective gene cluster (H1-H5). Samples are annotated based on their assigned dominant-species group.

Interestingly, metagenomes that were dominated by *B. breve* or *BL. longum* also contained genes from the H5 cluster, emphasizing their ability to utilize HMOs based on lacto-N-biose (LNB)^16^. However, samples from the “Mixed” groups exhibited minimal or no HUGs, suggesting either alternative genes for HMO utilization or a lack of capacity to utilize HMOs altogether.

### *BL. infantis* does not colonize the infant gut in early breastfeeding weeks

*BL. infantis* is the most proficient HMO-utilizer in the infant gut^11, 19^, thus we expected that *BL. infantis* will have a fitness advantage in the breastfed infant gut from the initial days of breastfeeding. However, despite the majority of infants in our cohorts that were breastfed since birth, *BL. infantis* was primarily detected starting at 10 weeks of age (**Figure 6A**). Overall, *BL. infantis* exhibited the highest abundance at 10-25 weeks, followed by a gradual decrease in abundance (**Figure 6A**). To corroborate these findings, we examined additional infant cohorts from Sweden^9^ and the United States^41^, comprising samples from earlier and/or subsequent time points. In the Swedish cohort, the presence of *BL. infantis* was not observed at birth almost at all, reaching its peak prevalence at 4 months, followed by a gradual decline in both prevalence and relative abundance by 12 months (**Figure 6B**). In the US cohort, no infants had any *BL. infantis* in the first week of life, and only one out of 77 infants exhibited its presence in the second week of life (**Figure 6C**). When comparing the prevalence of *BL. infantis* across all cohorts, we found that overall, the Israeli cohort had the highest prevalence (with 30% of infants having a detectable amount of *BL. infantis* at any time point) compared to the Swedish and US (20% and 2%, respectively; **Figure 6D**). The differential prevalence of *BL. infantis* across cultures has been recently observed in additional geographic locations^29^, and our computational approach allows us to continue to study this intriguing phenomenon in already existing datasets.

**Figure 6:**
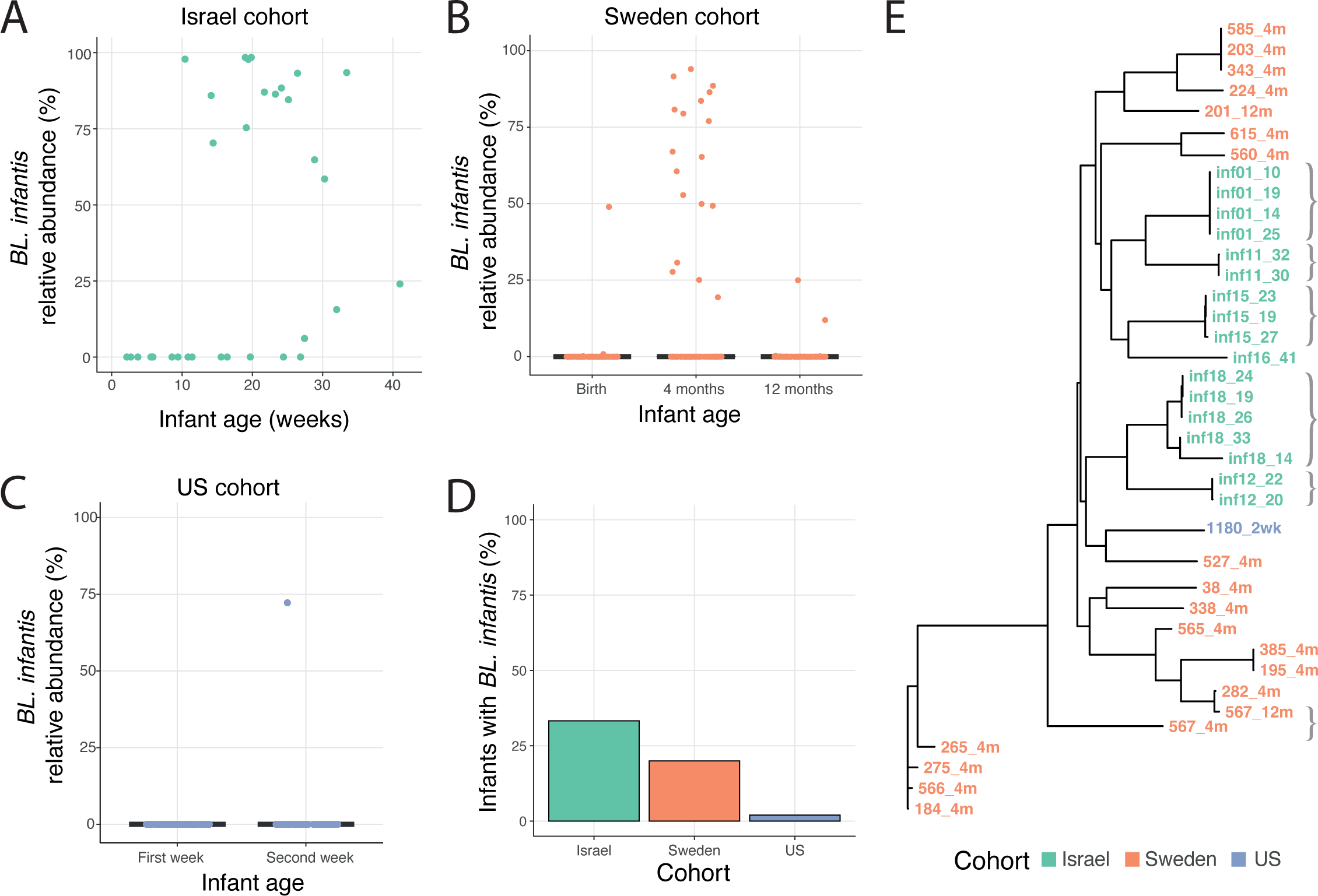
Late colonization and low prevalence of *BL. infantis* in the infant gut. **(A)** Relative abundance of *BL. infantis* in our Israeli cohort, among infants where *BL. infantis* was detected at any time point. **(B-C)** Relative abundance of *BL. infantis* at different time points in samples from **(B)** a Swedish cohort and **(C)** a US cohort. **(D)** Percentage of infants harboring *BL. infantis* at any time point in the Israeli, Swedish, and US cohorts. **(E)** Phylogenetic tree displaying *BL. infantis* strains found across all cohorts. The dominant strain from each sample is represented, and strains are color-coded based on their corresponding cohort. Brackets group strains from the same infant, emphasizing the stability of *BL. infantis* and the similarity of strains within each cohort.

To examine variations between *BL. infantis* strains across the different countries and to assess their stability, we next focused on *BL. infantis* strain-level composition. We found high similarity at the strain-level between samples from the same geographic location (**Figure 6E**). Within the Israeli cohort, infants that had detectable amounts of *BL. infantis* at multiple time points had nearly identical strains throughout our study (the exception being two time points of inf18; **Figure 6E**, gray brackets). However, in the single infant of the Swedish cohort that had *BL. infantis* at two time points (infant 567, 4 months and 12 months), different strains were observed, perhaps related to the large time difference between the samples (**Figure 6E**, bottom gray brackets). This observation suggests that different strains of *BL. infantis* are prevalent in different countries, and in the Israeli cohort, once *BL. infantis* has established in the infant gut, it is rarely displaced by other strains.

## Discussion

In this study, we introduced an innovative approach to quantify *BL. infantis* and distinguish it from *BL. longum* in metagenomic data. This method enables researchers to concentrate on studying this distinct subspecies and its associations with HMOs from existing metagenomic data. We employed our approach to explore the diversity within the infant gut microbiome and discover the lack of associations with HMOs present in mothers’ milk. Our analysis revealed that the variability between samples was greatly influenced by the dominant *Bifidobacterium* species in each sample.

Previous research has suggested that colonization of *Bifidobacterium* species in the infant gut may be influenced by priority effects^38^. However, our study revealed substantial changes in the dominant *Bifidobacterium* species within the same infant over the course of several weeks (**Figure 3A,B**). This indicates that over time there are additional factors responsible for *Bifidobacterium* species prosperity, such as species competition and cross-feeding. For example, it was reported that *B. breve*, despite having limited ability to utilize HMOs, can outcompete stronger competitors if introduced early into a microbial community^38^. In addition, *B. breve* has the capacity to cross-feed on monosaccharides derived from HMOs by other *Bifidobacterium* species^42, 43^. This implies that *B. breve* may initially dominate the population when carbohydrates are available, but subsequently loses the competition to other *Bifidobacterium* species once these carbohydrates are depleted. In our data we found that the *B. breve* dominated group had a diverse microbial population (**Figure 3C**), perhaps since it cross feeds on HMOs derivatives from other species. More research is needed to understand the microbial shifts in the infant gut and the effect cross-feeding has on the microbial dynamics in the infant gut.

While the HMOs present in a mother’s milk remained relatively stable over numerous weeks, we observed notable changes in the infant gut microbiome during this time period. This suggests that even subtle variations in the composition of breast milk may have an impact on the development of the gut microbiome. Alternatively, it is possible that other components present in breast milk, such as cytokines, microRNAs and antibodies, play a role in influencing the infant’s microbiome^44, 45^. Our findings indicate that there were no significant correlations between HMOs and *Bifidobacterium*, further supporting the idea that additional factors beyond HMOs are involved in shaping the infant’s microbiome. Previous studies^22–24, 26^ have examined the composition of the microbiome and its relationship with HMOs using 16S-rRNA amplicon sequencing. These studies have reported varying results, with some finding no significant correlations, while others observed modest correlations. Interestingly, some of the studies identified a negative correlation between *Bifidobacterium* OTUs and multiple HMOs^22, 24^. This could be attributed to a decrease in the *Bifidobacterium* population within the infant gut over time, coupled with an increase in specific HMOs, in line with our findings regarding LSTc and *BL. Infantis* (**Figure 4B**). Additionally, it is possible that such specific correlations may be observable only using a larger cohort.

Finally, we revealed that *BL. infantis* does not colonize the infant gut in the early weeks of breastfeeding and exhibits a relatively low prevalence. Previous studies have reported a low prevalence of *BL. infantis* during early time points^29, 35^, however these studies did not incorporate frequent sampling in the first months of life, thus failing to precisely determine the timing of *BL. infantis* arrival. Taft et al.^29^ proposed that the arrival of *BL. infantis* is influenced by the breastfeeding practices of a given country^29^. Countries with lower breastfeeding rates are likely to have a lower prevalence of *BL. infantis*, resulting in infants acquiring *BL. infantis* at a later stage through horizontal transfer. In our cohort, the prevalence of *BL. infantis* was lower compared to other *Bifidobacterium* species, yet it exceeded the prevalence observed in cohorts from Sweden and the United States. This prevalence is relatively high compared to other large cohorts from Western countries, such as Finland^46^, US^29^ and UK^47^. Further research, especially longitudinal sampling of infants and their surroundings, is required to elucidate the timing and sources from which infants acquire *BL. infantis* and to comprehend the differences observed between countries.

## Methods

### Sample collection

Breast milk and stool samples were collected as part of the Breast Milk Baby (BMB) cohort from mothers and infants from birth till one year old. Stool samples were collected using eSwab® with 1 ml of liquid Amies medium + 1 regular FLOQSwabs® (Copan) in order to preserve bacterial population. Breast milk samples were collected by pump or manually and stored in sterile tubes. Both sample types were collected by mothers in their homes and stored at 4°C for up to 24 hours and then shipped to the lab and stored long term at -80°C. All mothers have agreed to participate in our study, which was approved by the Hebrew University’s Institutional Review Board (IRB, approval number 20042021), and signed our consent forms.

### Metagenomic library construction and sequencing

DNA was extracted from stool samples using DNeasy PowerSoil Pro Kit (QIAGEN). Illumina sequencing libraries were prepared using Nextera XT DNA Library Preparation kit (Illumina) according to the manufacturer’s recommended protocol with half the volume and DNA. Samples were sequenced using Illumina single-end 150bp sequencing on a NextSeq 500 device.

### *B. longum* subspecies quantification

*B. longum* subspecies specific markers were found using 118 *B. longum* reference genomes downloaded from the NCBI **(Supplementary Table 2)**. Reference genomes were classified to *B. longum* subsp. *longum* (*BL. longum*) and *B. longum* subsp. infantis (*BL. infantis*) based on NCBI annotation, so that there were 101 *BL. longum* and 17 *BL. infantis* genomes. Two *BL. infantis* genomes did not contain the H1 cluster^19^ and therefore were suspected as mis-annotated and were excluded from further analysis. PanPhlan3^48^ was used to find subspecies specific markers. Genes were considered subspecies specific if they were present in 90% of the relevant genomes and not present at all in the other subspecies. Marker genes were filtered to be species specific using Blastn 2.12.0. Genes that were found in another species and had >90% alignment and matched over 50% of the marker gene were filtered out. Final markers included 118 *BL. infantis* and 90 *BL. longum* markers **(Supplementary Table 3)**. The MetaPhlAn database was customized using described instructions (https://github.com/biobakery/MetaPhlAn/wiki/MetaPhlAn-4) and then MetaPhlAn was used with the customized database using --index and --bowtie2db with our customized database.

To verify the results, subspecies of *B. longum* were determined using qPCR with specific primers^49^ for *BL. longum* (F: GTGTGGATTACCTGCCTACC, R: GTCGCCAACCTTGACCACTT) and *BL. infantis* (F: ATGATGCGCTGCCACTGTTA, R: CGGTGAGCGTCAATGTATCT). The efficiency of the primers was assessed by testing them in five dilutions. qPCR was performed at 95°C for 10 seconds, followed by 40 cycles of 95°C for 10 seconds and 60°C for 30 seconds. The ratio between *BL. infantis* and *BL. longum* was calculated using the delta-delta Ct method.

To use our tialored MetaPhlAn database see our GitHub page (https://github.com/yassourlab/MetaPhlAn-B.infantis/)

### Metagenomic analysis

Host reads were removed using an in house pipeline by aligning reads to the human genome by Bowtie2^50^. Samples were filtered and trimmed for Nextera adaptors using fastq-mcf, ea-utils^51^. Taxonomic profiling was done using MetaPhlAn4^33^ as described above. Functional profiling was done using HUMAnN^48^. HUGs^40^ were selected from previously described HMO clusters^32^. Strain analysis was performed using StrainPhlAn 4^33^. Further analysis was done using an in house R (4.2.2) script utilizing dplyr^52^ (1.1.2), tidyr^53^ (1.3.0), tidyverse^54^ (2.0.0). Plots were created using ggplot2^55^ (3.4.2) and ggforce^56^ (0.4.1), colors were used from RColorBrewer^57^ (1.1-3) and pals^58^ (1.7). Heatmaps were created using pheatmap^59^. Alpha and beta diversity were calculated using the vegan^60^ (2.6-4) package and the PcoA was created using the ape^61^ (5.7-1) package. Phylogenetic tree was produced using ggtree^62^ and sankey plots were made using ggsankey. Additional cohorts were downloaded from NCBI Sequence Read Archive using BioProject PRJEB6456 and PRJNA591079, for the Swedish^9^ and US^41^ cohort respectively.

### HMO quantification

HMO standards used in this study were purchased from Dextra Laboratories, United Kingdom. These included 2′-fucosyllactose (2′FL), 3-fucosyllactose (3’FL), 3′-sialyllactose (3′SL), 6′-sialyllactose (6′SL), lacto-N-tetraose (LNT), disialyllacto-N-tetraose (DSLNT), Lactodifucotetraose (LDFT), lacto-N-difucohexaose 1 (LNDFH), lacto-N-fucopentaose (LNFP) 1, 2, and 3, lacto-N-hexaose (LNH), lacto-N-neotetraose (LNnT) and sialyl-lacto-N-tetraose (LST) a, b and c. Linear B6-Trisaccharide was used as an internal standard.

HMO quantification was performed as previously described^63, 64^. Briefly 5 µl of human milk was combined with Linear B-6 Trisaccharide (Dextra Laboratories, UK) and HPLC grade water, then subjected to C18 columns (Thermo Scientific #60108-390) and carbograph columns (Thermo Scientific #60302-606) to remove proteins and salts respectively. Samples were labeled using 2-aminobenzamide (2-AB, Sigma) for 2 hours at 65°C. Excess 2-AB was removed using Silica columns (Thermo Scientific, #60300-482). Samples were separated by HPLC with fluorescence detection on a TSKgel Amide-80 column (Tosoh Bioscience, Tokyo, Japan) with a linear gradient of a 50 mM ammonium formate/acetonitrile solvent system. Retention times of purchased standard HMOs were used to annotate HPLC peaks. 6’SL and LNT peaks could not be separated and therefore, were calculated together. The amount of each individual HMO was calculated based on normalization to the internal standard. **(Supplementary Table 4)**. The relative abundance of each of the individual HMOs was determined by setting the sum of the 16 identified oligosaccharides as 100% total HMOs.

## Data availability

Human-filtered metagenomic sequencing data was deposited in SRA under BioProject PRJNA994433.

## Acknowledgements

This work was funded in part by the Azrieli Foundation grant for faculty fellows, by the Israel Science Foundation grant 2660/18 and by the Waterloo foundation.

## Author contributions

D.E. established the cohort, generated the sequencing data, quantified the HMO abundance and performed all the analysis. S.S. assisted with the experimental setup. E.J.K. guided and taught the HMO quantification method. M.Y. guided the work. D.E. and M.Y. wrote the manuscript.

## Competing interests

The authors declare no competing interests.

---

**Supplementary Figure 1:**
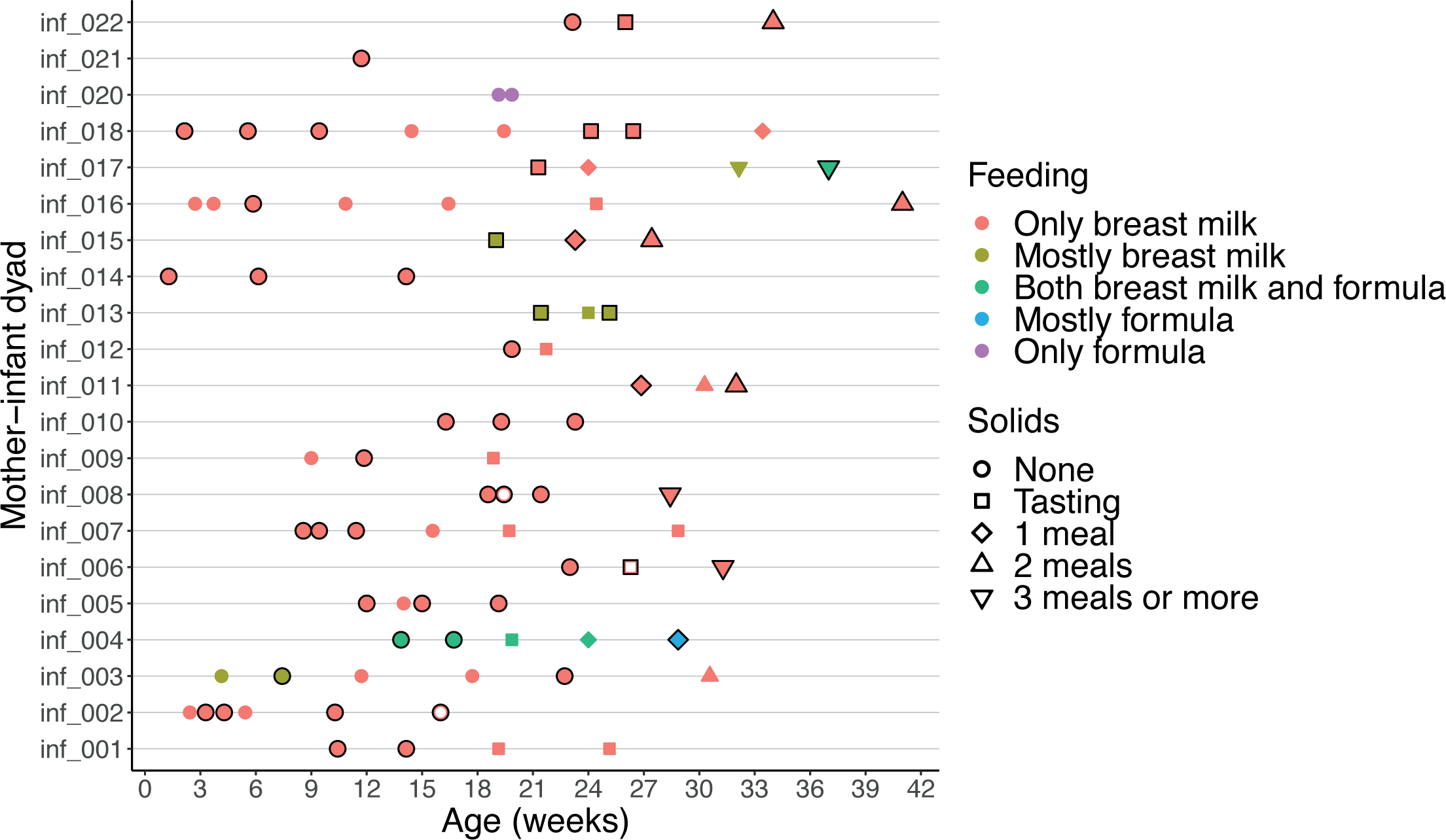
Cohort Overview. Overview of all samples collected as part of the cohort. Each data point represents a pair of an infant stool sample and a mother’s breast milk sample. Data points with a black border indicate breast milk samples used for HMO quantification, while filled-in data points represent stool samples subjected to metagenomic sequencing. The color of the points represents the feeding practice, and the shape of the points indicates the amount of solids consumed by the infant per day.

**Supplementary Figure 2:**
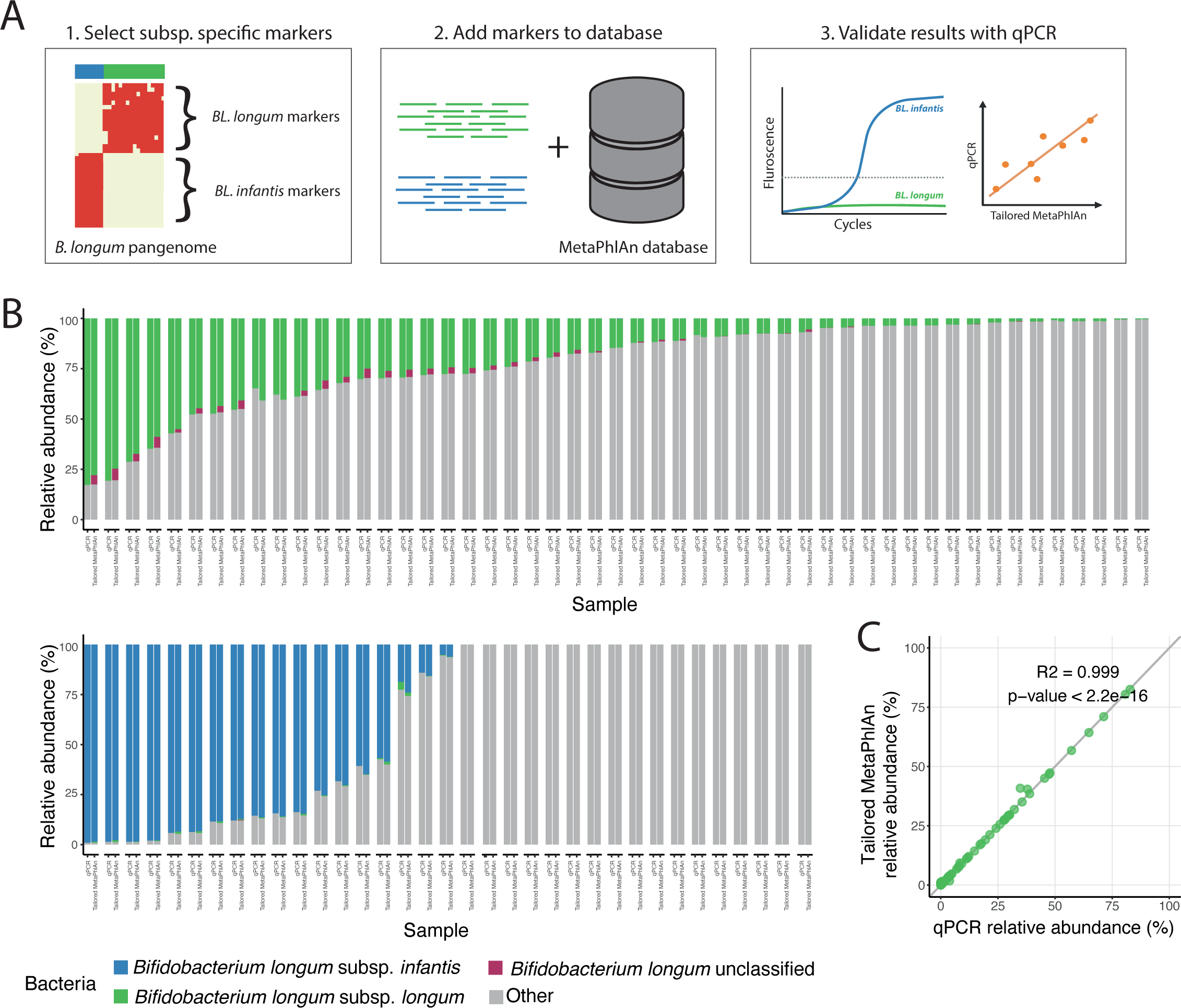
B. longum subspecies specific marker genes. (A) Illustration depicting the process of selecting and validating B. longum subspecies marker genes and their addition to the MetaPhlAn database. (B) Relative abundance of B. longum subspecies using qPCR and the customized MetaPhlAn method. Each sample is represented by two bars, one for each method. B. longum that was not assigned to either subspecies by our tailored MetaPhlAn method was classified as B. longum unclassified (burgundy). (C) Validation of the computational approach, comparing the tailored MetaPhlAn marker-gene quantification results to the experimental qPCR results for BL. longum. The quantification using MetaPhlAn includes bacteria assigned as B. longum unclassified.

**Supplementary Figure 3:**
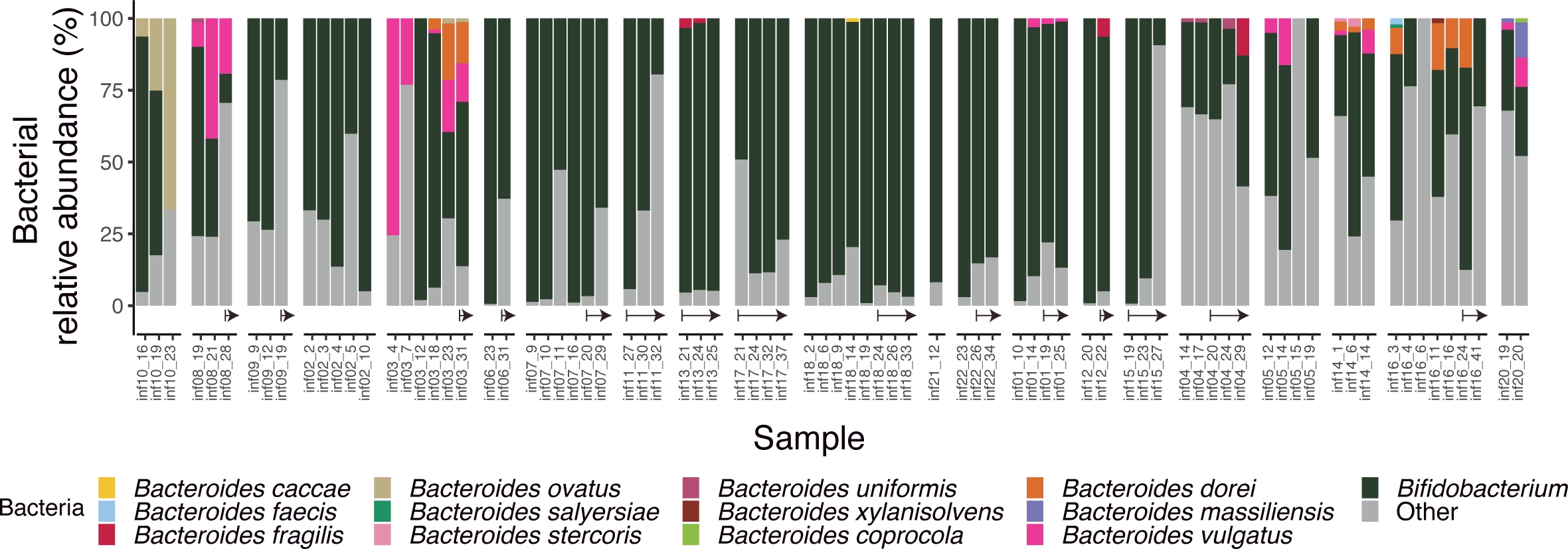
Bacterial composition of the infant gut. Microbial composition in all samples highlighting the Bifidobacterium genus and Bacteroides species. All other bacteria are classified as “other” (gray). Arrows indicate samples taken after the introduction of solid food.

**Supplementary Figure 4:**
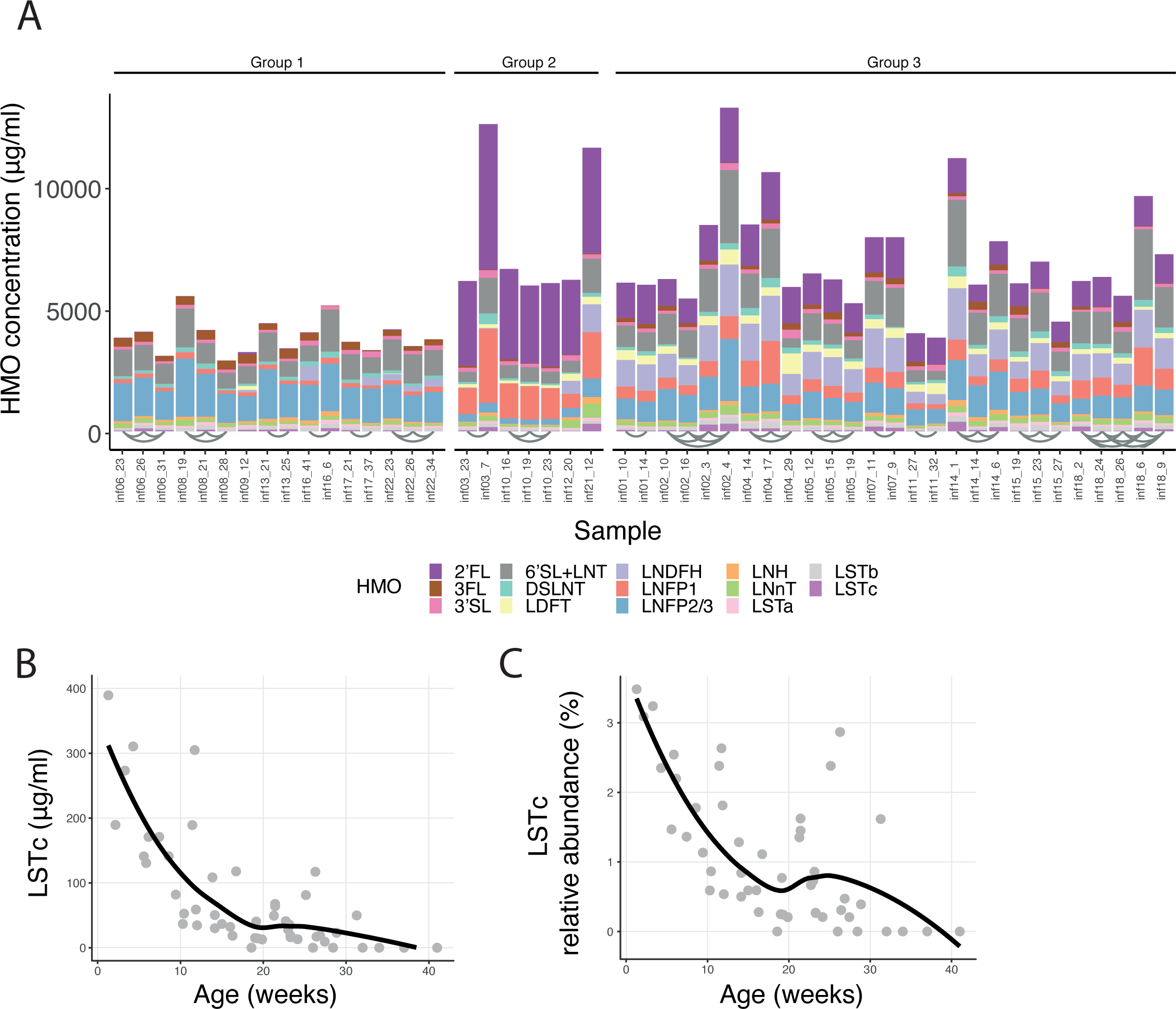
HMO composition in mothers’ milk. (A) Absolute concentration of 16 HMOs measured in mothers’ milk. Samples are categorized into three groups based on their HMO profiles, and arches connect samples obtained from the same mother. (B-C) LSTc (B) absolute concentration and (C) relative abundance over time in all breast milk samples measured. The black line represents the pattern.

**Supplementary Figure 5:**
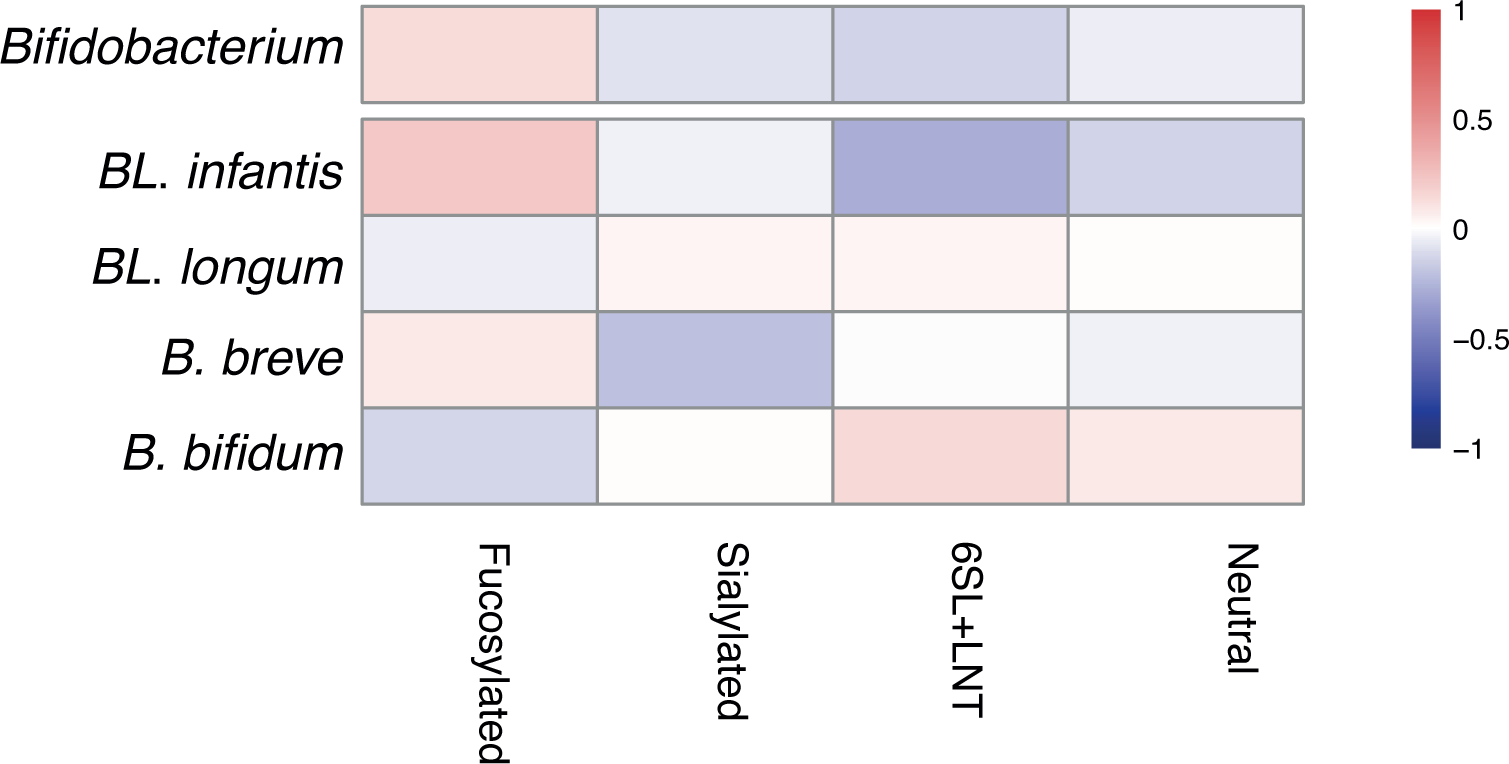
Correlation of HMO types with the infant gut microbiome. Pearson correlations between HMO types (neutral, fucosylated and sialylated) and the Bifidobacterium genus, as well as the main individual species and subspecies (BL. infantis, BL. longum, B. breve and B. bifidum). None of these correlations were found to be statistically significant.

## Notes

https://github.com/yassourlab/MetaPhlAn-B.infantis/

### Competing Interest Statement

The authors have declared no competing interest.

